# MuSkeMo: Open-source software to construct, analyze, and visualize human and animal musculoskeletal models and movements in Blender

**DOI:** 10.1101/2024.12.10.627828

**Authors:** Pasha A. van Bijlert

## Abstract

Musculoskeletal models for multibody dynamic analysis provide unique insights into human and animal movement. Although some biomechanical simulators provide model-building tools, these presuppose substantial preprocessing by the user, and resulting models are generally not cross-platform compatible. Thus, the workflow from anatomical 3D scans to musculoskeletal model is time-consuming, requiring numerous processing and conversions steps between software packages, and the process differs between simulators. Despite the popularity of musculoskeletal modelling within biomechanics, no cross-platform, open source software package exists for constructing musculoskeletal models.

Here, I introduce MuSkeMo: A software suite for defining 3D musculoskeletal models entirely within Blender (open-source 3D computer graphics software). MuSkeMo provides a visual interface, enabling users to interactively define all aspects of a musculoskeletal model (including rigid bodies, skeletal geometry, joint centres, muscles, landmarks, and body-fixed reference frames). MuSkeMo can calculate 3D inertial tensors from arbitrary meshes (e.g., from CT scans), and also implements automated convex-hull based mass-estimation approaches from the literature. Joints can be defined using shape-fitting of bony surfaces, and muscles can wrap around primitive shapes.

Models can be analyzed within MuSkeMo using popular pose-sampling procedures, or exported to multiple text-based formats for use in biomechanical simulators. A conversion script to OpenSim is included.

MuSkeMo is compatible with models created for popular biomechanical simulators (OpenSim and Gaitsym). MuSkeMo can import these models and simulation trajectories, enabling users to create publication-ready stills and animations with Blender’s ray tracing. These visualisations include volumetric muscles based on the contractile parameters, which can be more visually intuitive than traditional constant-diameter tube segments.

Whether the end goal is a highly-detailed subject-specific human model, or a simplified animal model, MuSkeMo includes features that can aid this process. By consolidating many elements of common model construction workflows into a cohesive package, MuSkeMo substantially simplifies musculoskeletal modelling.

## Introduction

For most animals, the ability to move is essential to their survival, and impairments to the musculoskeletal sytstem can have far-reaching consequences. The study of animal movement is thus fundamental to many fields, including clinical and veterinary biomechanics, comparative anatomy, human orthopedics, evolutionary biology, sports-medicine, and paleontology [1–6]. Technological breakthroughs in experimental measurement techniques (e.g., cinematography) have paved the way for many insights into animal movement [7–13]. However, measurements of animals in motion generally have a limitation: On their own, they often provide limited insights into the internal mechanics of an organism, and measurements of internal stresses and strains require invasive surgical procedures [8, 11, 12, 14].

Musculoskeletal models—here defined as mathematical, linked-chain rigid body models with force elements representing the musculoskeletal system—are therefore invaluable in the study of animal movement (Fig 1). Combined with external measurements of kinematics or ground reaction forces, musculoskeletal models enable us to look inside a moving animal [15, 16]. Using models, it is possible to parametrize anatomy in ways that cannot be achieved through physical experiments. For example: studies on the effects of body size on running performance [17], alterations of musculotendon architecture on gait selection [18, 19], the instrinsic perturbation-resistance that muscles provide [20–22], and numerous neuromechanical studies quantifying different aspects of healthy and impaired gait [23–25]. Musculoskeletal models have also provided insight into the evolution of the musculoskeletal system of a variety of tetrapod animals [26–35].

**Fig 1.**
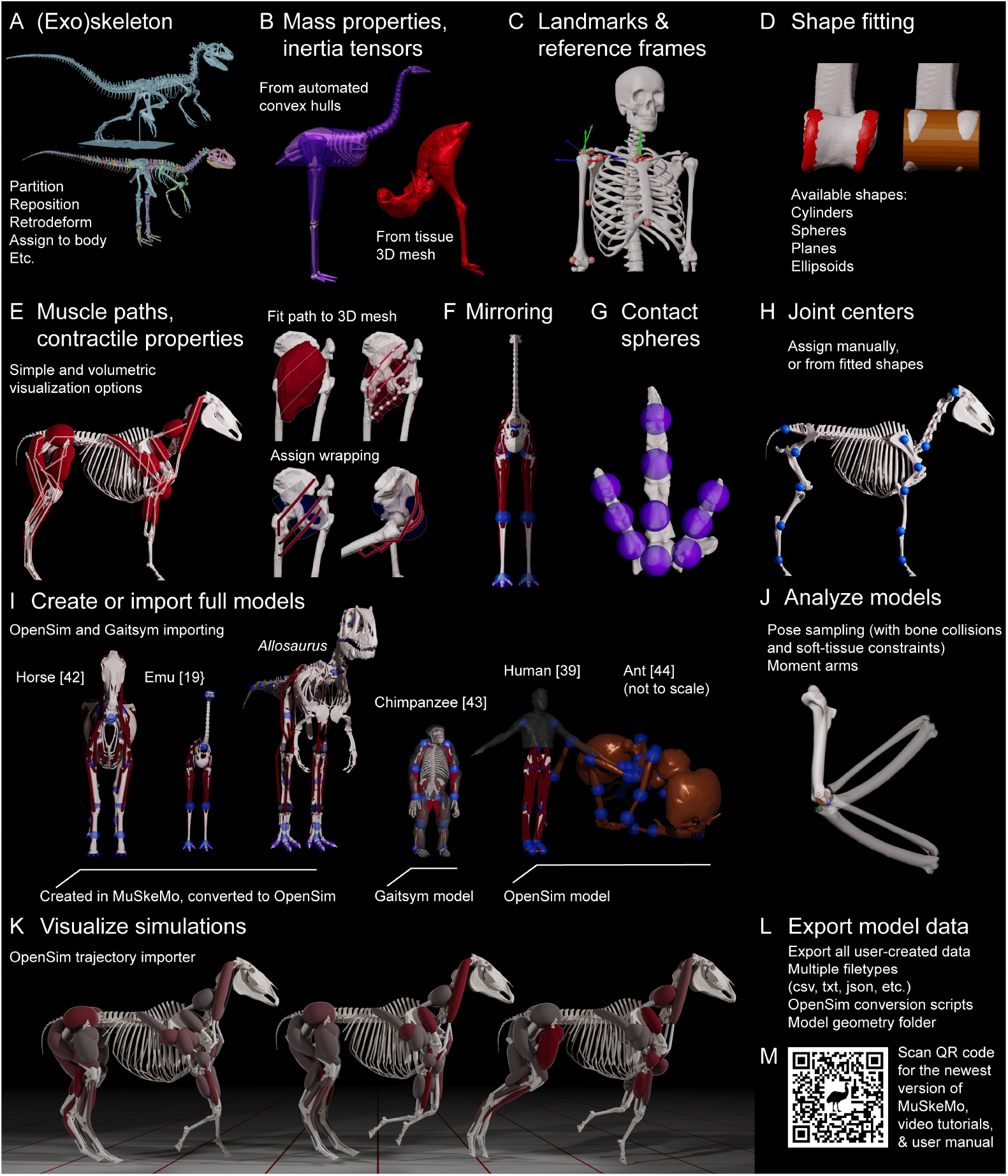
MuSkeMo consolidates common workflows in musculoskeletal model construction, analysis, and visualization into one cohesive package. A: Split meshes into relevant segments, which can be individually parented to rigid bodies. B: Estimate mass properties directly from tissue meshes, or using convex-hull based approaches from the literature. C: Add landmarks and reference frames. Human skeleton from [36], landmarks and frames following ISB standards [37]. D: MuSkeMo implements shape fitting of cylinders, spheres, ellipsoids, and planes. E: Define 3D muscle paths. MuSkeMo implements a path fitter for estimating lines of action from 3D meshes of muscles, and implements cylinder wrapping from [38]. Human muscle and bone meshes from [39]. F: MuSkeMo provides mirroring functions for symmetric model creation. G: Assign contact sphere locations. H: Joint centers can be defined manually, or using fitted shapes. I: MuSkeMo can be used to create new models or modify existing ones, using the OpenSim [40] and Gaitsym [41] importers. Horse model from [42], emu from [19], *Allosaurus* is unpublished, chimpanzee Gaitsym model from [43], human from [39], and ant from [44]. J: MuSkeMo implements soft-tissue constrained pose sampling analyses, and moment arm analyses. Pigeon data from [45]. K: While MuSkeMo is not a simulator, it is possible to visualize simulations using the trajectory importer. Modified from [42]. L: All model data can be exported in world and local coordinates as .CSV or other text files. OpenSim conversion scripts are provided. M: MuSkeMo is available from:

Advances in digital imaging and measurement techniques over the last three decades have made it possible to acquire both anatomical and motion data in unprecedented three-dimensional detail [13, 46–54]. This has motivated an increased interest in incorporating 3D data in human and animal models and simulations (e.g., [4, 33, 35, 36, 40, 41, 43–45, 55–60]).

Fully defining 3D musculoskeletal models often requires a large number of steps, that most analyses thus have in common [39, 40, 61, 62]. These include determination of inertial properties, computing global and relative positions and orientations of model components, and defining the relative hierarchies of these components (i.e., construction of a “digital marionette” [62]). Often, more components are added to the model to define its dynamic behaviour in simulations (e.g., muscle paths, wrapping surfaces, geometry for contact interactions) or for visualization (e.g., bone meshes). Despite these construction steps being shared by most musculoskeletal analyses, and the increasing popularity of these approaches, relatively few open source toolkits to construct biomechanical models exist. Several model building workflows exist for OpenSim: NMSBuilder [63], OpenSimCreator [64], and certain tools enhance the modelling capabilities of OpenSim (e.g., [65]). Gaitsym has a built-in model construction workflow [66]. Other model builders exist (briefly reviewed in [65]) that are not open-source and free, so will not be dealt with here. Apart from being designed for specific simulators and thus not generalizable to other software, the aforementioned approaches generally assume that most or all of the 3D mesh-processing steps have already been performed in other 3D computer-assisted design (CAD) software. This has motivated certain groups to develop custom workflows using various combinations of open and closed-source programs (e.g., [29, 34, 62, 67, 68]). Other groups have proposed methods to estimate certain input parameters that are used by musculoskeletal models (e.g., convex hull approaches for estimating inertial properties of body segments [69–71]), but these are not currently implemented in existing model builders. An open-source, simulator-agnostic toolkit for defining musculoskeletal models could therefore benefit many researchers in the biomechanics community.

Here, I introduce MuSkeMo: A suite for processing, analyzing, and visualizing 3D biological scan data for biomechanical and morphological analyses. MuSkeMo functions as a plugin for Blender (blender.org), open-source 3D modelling program. MuSkeMo is richly featured, enabling the user to fully-define musculoskeletal models. Starting from 3D meshes (e.g., from biological scans), MuSkeMo enables interactive model construction through its graphical user interface (GUI), while maintaining extensibility with custom Python scripts for advanced users. MuSkeMo has the following features:

- Determination of 3D inertial tensors from triangulated meshes (Fig 1B)
- Implementation of several convex-hull based mass-estimation approaches from the literature [69, 71] (Fig 1B)
- Aggregating meshes different inertial properties into a single rigid body using the parallel axes theorem
- 3D Landmarking and defining local and anatomical reference frames (Fig 1C)
- Fitting of geometric primitives to bony (articular) surfaces (Fig 1D)
- Defining path-point muscles, including simple wrapping operations (Fig 1E). Muscle paths can be fitted from 3D models acquired from CT/MRI segmentations
- Mirroring scripts for bilaterally symmetric models (Fig 1F)
- Assigning contact sphere locations (Fig 1G)
- Defining joints (manually, or via geometric primitive shape fitting) (Fig 1H)
- Automated joint-pose sampling scripts that check for mesh intersections (Fig 1M)
- Analyses of musculotendon (or ligament) lengths and moment arms
- Data export in customizable text-based formats (CSV, TXT, JSON, BAT, etc.), which can be reimported to combine different model iterations
- Compatibility with popular biomechanical simulators, including OpenSim and Gaitsym (Fig 1I)
- Publication-quality stills and animations, facilitated by a simulation trajectory importer (Fig 1K)
- A custom approach for representing the volume encoded in a Hill-type path point muscle, for more intuitive visual communication (compare Fig 1I&K)

By implementing MuSkeMo in Blender, Blender’s expansive and specialized 3D modelling functionalities are also available during the construction process. This provides a uniquely diverse toolkit during model construction, and consolidates all the necessary processing steps into one single program, preventing unnecessary conversion steps and possible loss of data. Because Blender is both powerful and open source, it has been gaining popularity in scientific fields that use 3D data. Of particular relevance to MuSkeMo are Myogenerator [72], which is geared towards Finite Element Analysis, and the Tetrapod Toolkit [73] and the Blender XROMM Toolkit [74], focused on kinematic analyses.

With its focus on musculoskeletal modelling, MuSkeMo provides a unique contribution to this thriving ecosystem of open-source research software in Blender. In the next sections, I will first describe how MuSkeMo has been implemented, highlight some of the biological research that has already used MuSkeMo in a variety of ways, and finally provide some examples in how MuSkeMo can synergize with existing scientific software in Blender.

## Design and implementation

### MuSkeMo’s design philosophy

MuSkeMo is primarily intended to be a comprehensive toolkit for defining, analyzing, and visualizing biomechanical data and musculoskeletal models. MuSkeMo achieves this by enabling the user to perform a wide array of transformations to translate biological 3D data to useful biomechanical information. The toolkit is designed to be species and scale invariant: It is equally straightforward to work on an ant or a dinosaur in MuSkeMo (Fig 1), as long as either high-quality 3D scans of the (exo)skeleton and other (soft) tissues are available, or fully-defined models of the species in question already exist. MuSkeMo is also intended to be simple to use via a graphical user interface (Fig 2), but extensible for experienced users via the Python application programming interface (API). Models created or modified in MuSkeMo can be exported for use in a simulator of choice. MuSkeMo is explicitly intended to be simulator agnostic—The user interactively defines a physical system, and should in principle be free to simulate it in their simulator of choice. By providing export capabilities for all components in a variety of text-based formats (e.g., CSV), MuSkeMo aims to maximize compatibility with existing simulators. Delivering a comprehensive featureset while remaining simulator agnostic is a challenge. Simulators can differ in their implementations of the same feature (e.g., muscle wrapping [75]), or simply omit them altogether. MuSkeMo prioritizes cross-platform compatibility, if models require simulator-specific features to be added, it is up to the user to add it in their simulator of choice.

**Fig 2.**
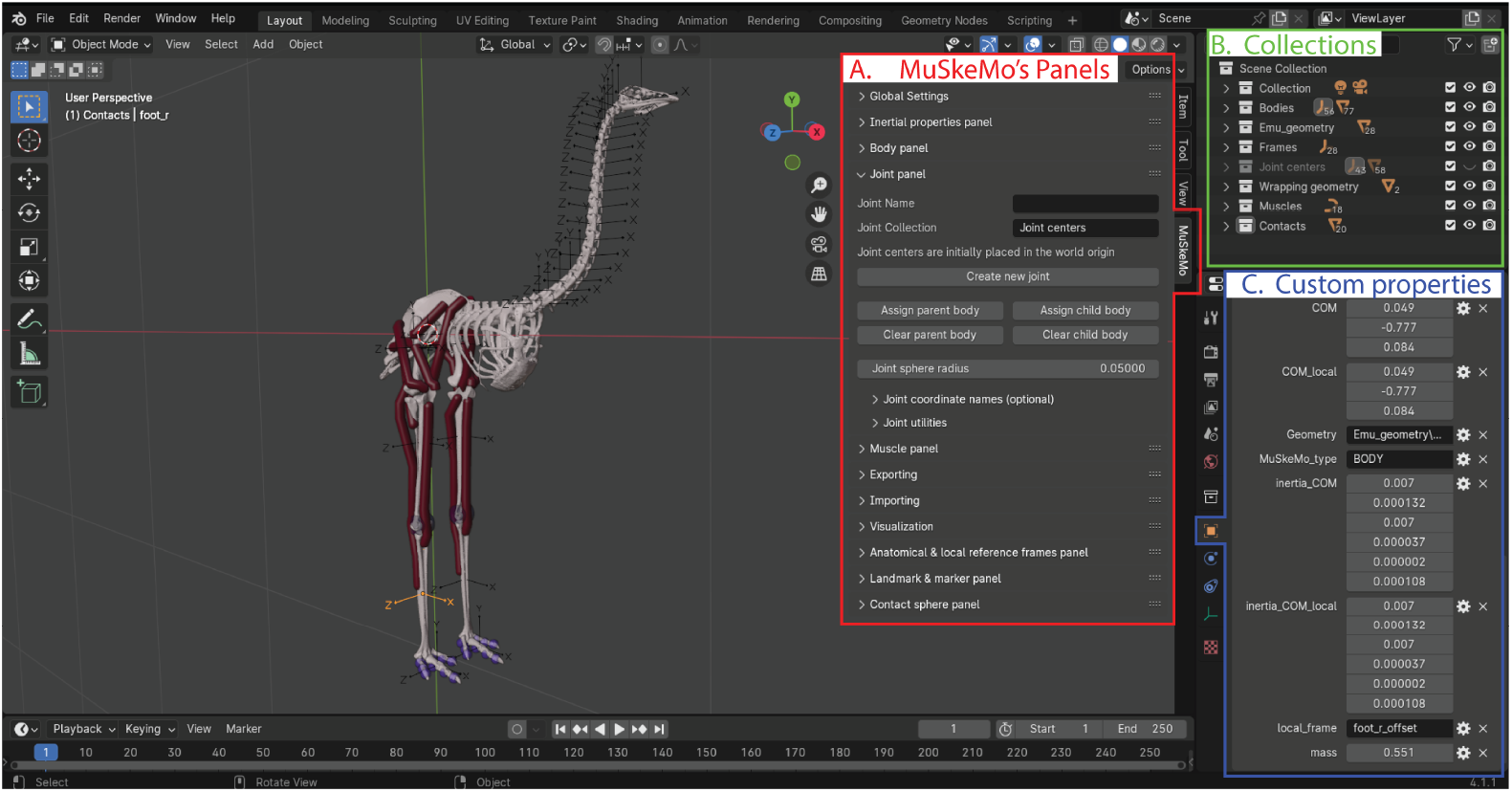
MuSkeMo provides a graphical user interface for biomechanical analysis and model construction in Blender. A: MuSkeMo’s features are accessed via its panels. B: MuSkeMo sorts model components in Blender “collections”, which act as folders or visibility layers. C: MuSkeMo stores component (meta)data in Blender’s “Custom properties”. Pictured are the data for the “foot r” BODY, which is selected in the model (the highlighted axes).

### MuSkeMo’s implementation

#### Dependencies and prerequisites

MuSkeMo is an add-on for Blender (tested for versions 4.0-4.2), and all of its main features have no other dependencies. Users who do not have adminstrator privileges to install Blender can download the “portable” version, which does not require installation before running the program. MuSkeMo is constructed using Blender’s Python API. All of MuSkeMo’s features can be accessed via the panels and buttons in the GUI (Fig 2A), and a detailed user manual describes all of its functionality and provides installation instructions (S1 File).

#### Model components, properties, and hierarchy

Model components are stored by MuSkeMo in named Blender “Collections” (Fig 2B). These act like folders or visibility layers, grouping similar components together. The different types of model components (e.g., rigid bodies and muscles), are abstract concepts that do not natively exist in Blender. MuSkeMo creates these by creating objects that do exist within Blender (e.g., arrows for rigid bodies, curves for muscles), and assigns extra metadata for each object type using Blender’s “Custom properties” system (Fig 2C). Each component is always assigned a “MuSkeMo type” that defines what type of component it is (e.g., “BODY” or “MUSCLE”), and a number of other MuSkeMo type-dependent inputs (e.g., “mass”). As the user progresses through the model-building steps, MuSkeMo’s defines the inter-relationships between these objects, and populates their inputs. The user manual (S1 File) provides a full list of available MuSkeMo types and corresponding inputs.

Blender keeps track of the positions, orientations, and scales of objects via their “matrix world”, which is a 4×4 transformation matrix [76]. MuSkeMo uses this matrix world (position, and orientation when relevant) internally to compute model properties with respect to the global reference frame. Users can define a local (anatomical) reference frame and assign this to a rigid body, in which case MuSkeMo computes all the transformations with respect to the local frame [76, 77]. MuSkeMo computes this as both Euler angles (using an XYZ, body-fixed decomposition order), and unit quaternions, implemented following [76–79] (see the user manual, S1 File for details).

Although it is possible to use Blender’s built-in rigging system for defining models (e.g., [73]), MuSkeMo uses Blender’s parenting system, because it provides more flexibility for supporting concepts such as local reference frames. The parenting system ensures that all the “children” follow the transformations of the “parent” object, essentially forming kinematic constraints. Under the hood, MuSkeMo makes use of this parenting system in Blender when defining inter-relationships between model components. However, because MuSkeMo also needs to compute local transformations during parenting, MuSkeMo provides its own functions that both call the Blender parenting system, and keep track of all the model components that have properties that need to be updated. These data are exposed to the user via custom properties (Fig 2C), and are also included during export.

#### Muscles, ligaments, and wrapping

Unlike all other model components in MuSkeMo, muscles are not parented in the aforementioned manner, because it is not possible to parent individual curve points to objects in Blender. Instead, this is achieved using the “modifier” system in Blender. Individual curve points (i.e., individual muscle path points) are attached to rigid bodies using a “hook modifier” for each point. Muscles have two main visualization options:

- A standard, equal radius “tube” style that precisely demonstrates the path, but ignores muscle volumes (Fig 2).
- A volumetric visualization style unique to MuSkeMo (Fig 1K). This visualization style accurately represents muscle volumes, but resamples the path to achieve this.

Both visualization styles are achieved by adding a “Geometry nodes” modifier to the muscles, and are customizable by the user. See the manual (S1 File) for details. Ligaments currently do not have a distinct implementation within MuSkeMo, but can simply be modelled as if they are muscles. Different simulators have different implementations of wrapping, or simply do not implement it at all [75, 80, 81]. MuSkeMo includes cylinder wrapping following [38], implemented using one Geometry nodes modifier per object (see S1 File).

#### Geometric primitive shape fitting

MuSkeMo implements geometric shape-fitting algorithms (Fig 1D), a common procedure to determine joint centers of rotation [62]. MuSkeMo can fit spheres [82, 83], ellipsoids [84], cylinders [85], and planes. The shape fitters are easily accessible via MuSkeMo’s GUI, and it is possible to match joint transformations to the resultant shapes, see the manual (S1 File).

#### Utility functions

The MuSkeMo download package includes a folder (“MuSkeMo utilities”) with several convenience functions. Currently, this includes the following scripts:

- MuSkeMo includes a Matlab (Mathworks) script that converts MuSkeMo-exported CSV files into an OpenSim file, using OpenSim’s Matlab API.
- MuSkeMoprovides a path fitting script that allows the user to estimate a line of action from a 3D mesh of a muscle (Fig 1E), similar to the Maya (Autodesk) implementation described in [34]. The fitting resolution can be specified by the user.
- Pose sampling is a popular technique for studying fossils [86, 87], thus far generally implemented in Maya. The MuSkeMo utilities folder includes a Python script that can perform a pose sampling analysis on a MuSkeMo model in Blender (Fig 1J). This script samples user-defined poses, checks for bone collisions/intersections (using Blender’s computationally efficient Bounding Volume Hierarchy system, implemented as “BVHTree”), and reports the corresponding length of a muscle or ligament for a given pose. These data are exported as a CSV file. This script can serve as a template for larger analyses such as [86, 87].
- A Python script that can be used to compute the minimum distance between two meshes, useful when articulating skeletal meshes or analyzing imported trajectories.

As more conversion and utility scripts become available (see Planned additions and possible community contributions), these will be included in the folder.

## Results

### Case studies and applications

This section presents some ways that MuSkeMo has already been used in the literature, highlights some projects that are still underway, and highlights potential use cases and workflows that MuSkeMo could facilitate.

#### Models for fundamental biomechanical research

MuSkeMo has been used to answer fundamental questions in the field of evolutionary biomechanics. For example: Van Bijlert et al. [19] recently constructed a generalized model of the emu using (a pre-release version of) MuSkeMo (Fig 3A). This model was used for predictive gait simulations in OpenSim Moco [40, 88]. They systematically varied the musculotendon parameters in their model, to investigate how different postural preferences affected optimal running styles. After validating the simulation outputs against empirical data, they were able to link a preference for crouched postures to the optimality of grounded running, resolving a long-standing question regarding bird running styles [89, 90].

**Fig 3.**
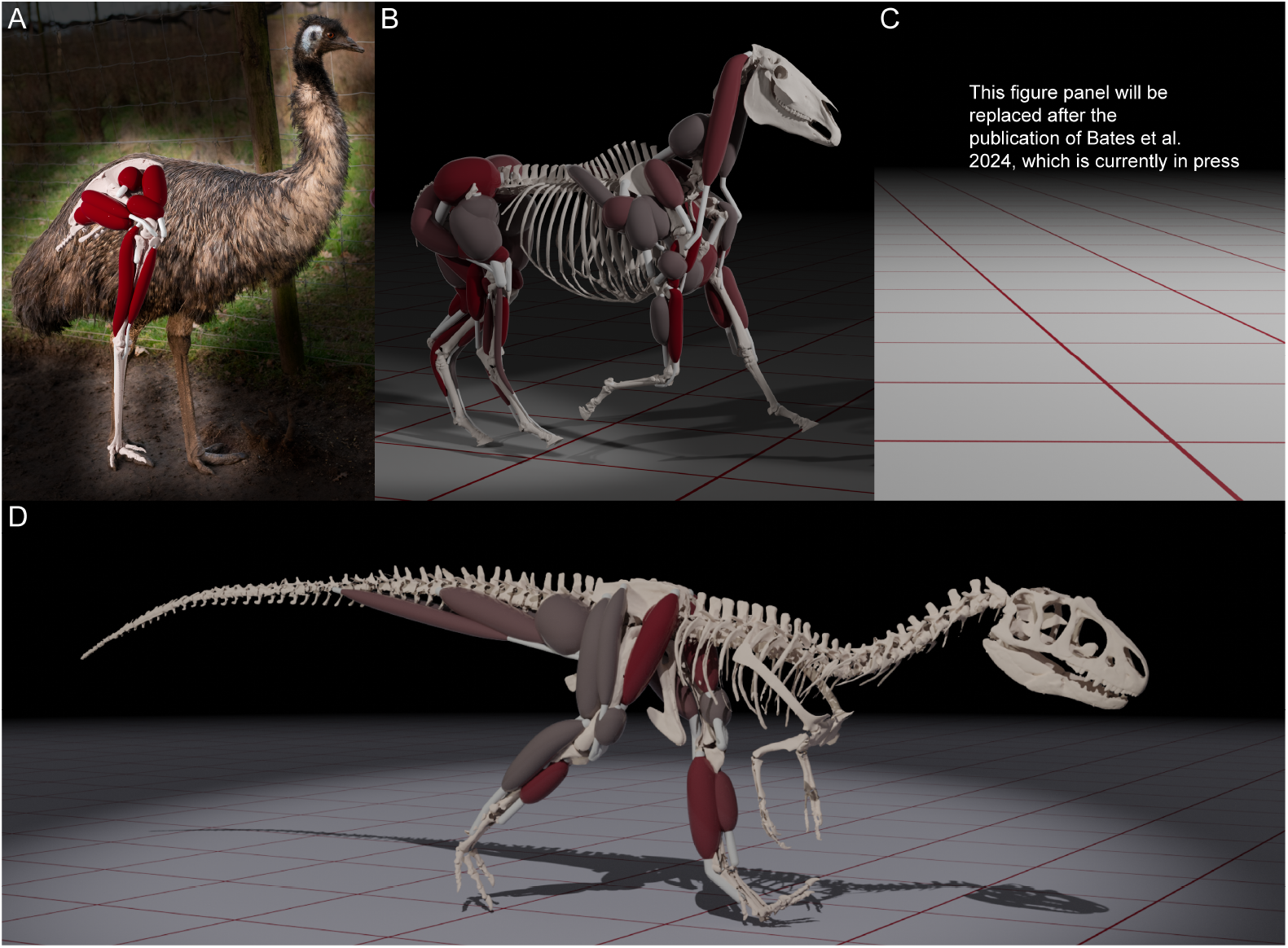
Examples of models either built or visualized using MuSkeMo. A: Emu model from [19], built using MuSkeMo, validated and simulated using OpenSim Moco. B: Horse model from [42], used for predictive and tracking simulations in OpenSim Moco. Used as the cover image of Volume 64, Issue, (2023) of Integrative & Comparative Biology. C: The *Australopithecus afarensis* simulations from [93] have been removed from this figure pending that paper’s official publication on December 18th 2024. D: *Allosaurus jimmadseni* model, developed for the BBC documentary “Secrets of the Jurassic Dinosaurs” (2023).

If adequate 3D scans are available, MuSkeMo can be used to construct models of extinct animals. For example, several groups are using MuSkeMo for analyses of non-avian dinosaurs [91, 92].

#### Models for (veterinary) clinical practice

A promising application of musculoskeletal models is the clinical practice, both in human and veterinary contexts [3, 40, 42, 94]. Van Bijlert et al. [42] presented a model of the horse, constructed in MuSkeMo Fig 3B). They simulated this model both in OpenSim using Moco [40, 88], and in SCONE using Hyfydy [56, 81]. Animal models can potentially play a valuable role in veterinary diagnostics, since non-human animals cannot verbally communicate pain. Personalized, subject-specific models [39], and models that simulate the effects of a surgery [4], have been been used to both improve model predictions, and even surgical outcomes. MuSkeMo could potentially facilitate this process: Not only is it possible to directly construct a subject-specific model using scans, it is also possible to modify existing models using imported (MRI or CT-based) images as a visual reference. Fig 1E demonstrates a possible workflow using human data from [39] (Subject 01): Using the segmented 3D mesh of the gluteus maximus (subdivided into an upper, intermediate, and lower section), the MuSkeMo’s path fitter was used to generate curves personalized path points (Fig 1E). These then served as a visual aid when applying cylindrical wrapping to simplified path-point curves.

#### Models as tools for education and outreach

Musculoskeletal biomechanics can be a challenging topic for scientific communication, due to the many concepts from physics that are abstract to non-experts. Effective visualizations can measurably improve the effect of scientific communication [95], and MuSkeMo provides the user with many tools for customizable, high-quality scientific visualizations. MuSkeMo has been utilized in this regard:

- Bates et al. [93] investigated the evolution of human running performance using predictive gait simulations of *Australopithecus afarensis*. Simulations were performed in Gaitsym, and the authors produced a video using MuSkeMo where the model variants ran side by side to directly compare the gaits Fig 3C).
- The BBC Documentary “Secrets of the Jurassic Dinosaurs” (2023) featured predictive running simulations of the carnivorous dinosaur *Allosaurus jimmadseni* Fig 3D). The model was constructed in MuSkeMo, simulated in OpenSim, and then visualized using MuSkeMo.

### Possible synergies within Blender

Several academic software plugins for Blender exist that are geared towards biomechanical analysis, which provide complementary functions to MuSkeMo.

- The Myogenerator plugin introduced by Herbst et al. [72] provides a plugin that enables 3D volumetric reconstructions of muscles, and outline workflows for retrodeforming (warping) skeletal meshes of fossil organisms for construction models for finite element analyses. Reconstructed muscles from Myogenerator could be used as the input mesh for MuSkeMo’s path fitter, to define the path point muscle in a musculoskeletal model (see Utility functions).
- X-Ray Reconstruction of Moving Morphology (XROMM) provides unprecedented accuracy in kinematic analyses [13, 47]. The Blender XROMMM toolkit [74] makes it possible to import XROMM trajectories into Blender. This could be combined with MuSkeMo in a variety of ways. E.g., the user could construct models using the in vivo measured poses as a starting point, and if muscles and ligaments are added using MuSkeMo, it would be possible to estimate changes in lengths during measured movements.
- The Tetrapod Toolkit [73] is aimed at rigging 3D (skin outline) models of animals, to determine stride parameters. MuSkeMo could be used to compute the inertial properties from 3D tissue outlines. Similar to with the XROMM toolkit, if the user defines an underlying musculoskeletal model using MuSkeMo, muscle and ligament length changes can be computed during the studied movements.

## Availability and future directions

### Repository, manual, and video tutorials

It is recommended to download the most recent release of MuSkeMo.zip directly from Github:

https://github.com/PashavanBijlert/MuSkeMo/releases.

Updated ZIP releases automatically include a PDF of the most recent version of the user manual (S1 File), which is also available separately on Github. To familiarize the user with the functionalities and recommended workflows, users can view the video tutorial series:

https://youtube.com/playlist?list=PLfgxaucAWlEp5-cavvXmdrTIWYT_tgZYK&si=OETJH5lB7YP0knc5

More tutorials will be added with example workflows.

### Bug reports and feature requests

Bug reports and feature requests can be submitted in the issues section on Github: https://github.com/PashavanBijlert/MuSkeMo/issues.

### Planned additions and possible community contributions

MuSkeMo v1.0 was developed by a single person, and more ambitious goals can be reached through contributions from the biomechanics community. Contributions can be proposed via Github, or by directly contacting the author.

#### Landmarking

While MuSkeMo’s focus is on musculoskeletal modelling, its features are useful more generally for morphological analyses in the biological sciences. In particular, landmarks are often used in geometric morphometric analyses [96, 97]. MuSkeMo has already implemented landmarking in Blender, and could already be used to this end in simple analyses. Future additions will extend the current functionality, and possibly add support for semi-landmarks.

#### Wrapping

MuSkeMo currently supports single object wrapping of cylinders. Multiple wrapping objects per muscle are possible, but must be separated by a path point or they will not be true tangent-curve solutions. This is due to the inherent limitations of having to define wrapping logic using Blender’s Geometry nodes. To my knowledge, it is currently not possible to implement iterative root-finding methods within a Geometry node, so multi-object wrapping algorithms that rely on this [38] cannot be implemented. This situation may change if either Blender implements the possibility of creating a custom type of curve via the API (currently, MuSkeMo MUSCLEs are implemented using standard “POLY” curves), or if Blender implements something akin to a “scripted expression” node within Geometry nodes. Users who may need more extensive wrapping within MuSkeMo are encouraged to propose solutions that may circumvent aforementioned hurdles.

#### Conversion to popular simulators

MuSkeMo is intended as a cross-platform solution for constructing musculoskeletal models. To this end, MuSkeMo is able to export all user-created data as customizable text files (CSV, TXT, etc.), allowing easy inter-simulator operability. The latest version of MuSkeMo includes scripts for the conversion of these files to OpenSim models. Importing of OpenSim and Gaitsym models into MuSkeMo is also supported. Planned additions include conversion scripts for both importing and exporting models to other popular biomechanical simulators (e.g., [75, 81, 98]). Community contributions, especially from experts using those simulators, are welcome and would accelerate this goal.

### Supporting information

#### S1 File. MuSkeMo manual

This user manual provides installation instructions and detailed descriptions of how to use all the functions in MuSkeMo. This manual also outlines some suggested workflows, lists all data types and inputs that MuSkeMo can create, and provides some background information in the Appendices.

#### S2 File. MuSkeMo

This ZIP-archive contains v1.0 MuSkeMo, and a separate directory with some Utility Scripts. The ZIP-archive can be directly installed in Blender (see the manual S1 File for installation instructions). New versions of MuSkeMo will be regularly pushed to Github.

## Acknowledgments

I would like to thank Matt Dempsey, Max Herde, Lars D’Hondt, and Zuhayr Parkar for providing feedback on previous versions of MuSkeMo. The ellipsoid fit Python algorithm was provided courtesy of Mark Semple, based on an implmentation by Yuri Petrov. David Eberly made the cylinder fitting algorithm available as pseudocode on his website Geometrictools.com. I am grateful to Karl Bates for providing comments on this manuscript.

